# SSAM-lite: a light-weight web app for rapid analysis of spatially resolved transcriptomics data

**DOI:** 10.1101/2021.09.29.462194

**Authors:** Sebastian Tiesmeyer, Shashwat Sahay, Niklas Müller-Bötticher, Roland Eils, Sebastian D. Mackowiak, Naveed Ishaque

## Abstract

1

The combination of a cell’s transcriptional profile and location defines its function in a spatial context. Spatially resolved transcriptomics (SRT) has emerged as the assay of choice for characterizing cells in situ. SRT methods can resolve gene expression up to single-molecule resolution. A particular computational problem with single-molecule SRT methods is the correct aggregation of mRNA molecules into cells. Traditionally, aggregating mRNA molecules into cell-based features begins with the identification of cells via segmentation of the nucleus or the cell membrane. However, recently a number of cell-segmentation-free approaches have emerged. While these methods have been demonstrated to be more performant than segmentation-based approaches, they are still not easily accessible since they require specialized knowledge of programming languages and access to large computational resources. Here we present SSAM-lite, a tool that provides an easy-to-use graphical interface to perform rapid and segmentation-free cell-typing of SRT data in a web browser. SSAM-lite runs locally and does not require computational experts or specialized hardware. Analysis of a tissue slice of the mouse somatosensory cortex took less than a minute on a laptop with modest hardware. Parameters can interactively be optimized on small portions of the data before the entire tissue image is analyzed. A server version of SSAM-lite can be run completely offline using local infrastructure. Overall, SSAM-lite is portable, lightweight, and easy to use, thus enabling a broad audience to investigate and analyze single-molecule SRT data.

**Availability and Implementation:** SSAM-lite is an open-source browser-based web application with source code freely available on Github via https://github.com/HiDiHlabs/ssam-lite. Stable releases can be accessed via https://ssam-lite.bihealth.org and https://ssam-lite.netlify.app, and developmental releases can be accessed via https://dev--ssam-lite.netlify.app. The source code for a locally deployable server version, SSAM-lite-server, is available on GitHub via https://github.com/HiDiHlabs/ssam-lite-server. Both versions require a modern browser with JavaScript and WebGL support. Detailed user guides and documentation can be found at https://ssam-lite.readthedocs.io.

## 2 Introduction

The biological function of a cell is governed not only by its expression profile but also by its location (Lee, 2017). A cell’s spatial embedding defines its cellular neighborhood and determines how intercellular signaling operates to achieve higher-order tissue function. Spatially resolved transcriptomics (SRT) has emerged as the assay of choice for characterizing cells in a tissue context (Burgess, 2019; Marx, 2021). There are a number of SRT methods, with each being able to resolve gene expression to various spatial resolutions, from anatomical features up to sub-cellular resolution of identifying single mRNA molecules (Asp et al., 2020). Single-molecule SRT methods usually require the assignment of each decoded mRNA spot to a cell, which first requires the cell to be identified via segmentation. Cell segmentation is usually performed by identifying cell landmark features such as the cell nucleus or protoplasm via DAPI or total mRNA density (Najman and Schmitt, 1994; Chen et al., 2015; Eng et al., 2019). However, accurate cell segmentation remains difficult due to many factors such as staining not covering all features of a cell, imaging artifacts, and overlapping cells (Thomas and John, 2017). Inaccurate cell segmentation can lead to misassignment of mRNA molecules to cells, leading to errors in downstream analysis such as misclassifying cell types. To overcome this issue, a number of computational tools have been developed to improve the assignment of mRNA molecules to cells (Qian et al., 2020; Prabhakaran et al., 2021), incorporate cell typing as part of the segmentation process (Littman et al., 2021), and perform cell-segmentation free analysis (Petukhov et al., 2020; He et al., 2021; Park et al., 2021). While these tools improve cell typing, they all share the problem of being specialized tools that require access to Linux command line terminals, programming expertise, and high-performance hardware. This renders them less accessible to a large proportion of the biomedical research community.

Our prior work (Park et al. 2021) demonstrated improved accuracy and sensitivity of spatial cell typing over traditional segmentation-based approaches by applying the SSAM algorithm to the mouse somatosensory cortex dataset profiled by osmFISH. In particular, our segmentation-free approach identified many more astrocyte cell types that were missed due to low signal. Furthermore, we could reconstruct the ventricle region that was missed due to high occlusion in the segmentation-based approach used in the original study of the data.

Here we present SSAM-lite which is an easy-to-use and lightweight browser-based web application on top of the segmentation-free SRT algorithm SSAM (Park et al., 2021) to make spatial cell typing accessible to biomedical researchers. SSAM-lite runs on modest hardware in any modern browser with JavaScript support and internet access, thus lowering the barrier to analyzing high-dimensional SRT data. To ensure privacy and security, data does not leave the user’s machine. Furthermore, our tool has an easy-to-use graphical user interface that provides intuitive visualizations of SRT data. SSAM-lite can be used on mobile devices to analyze smaller datasets. Departments or institutes with access restricted to local networks due to security reasons or which deal with extremely large datasets can make use of SSAM-lite-server. This is a server-side implementation of SSAM-lite that can be installed with minimal effort, providing offline access to SSAM-lite functionality and without limitations of client-side resources.

## 3 Methods

### 3.1 SSAM-lite

SSAM-lite builds on top of the SSAM algorithm (Park et al., 2021) (Figure 1). In brief, the algorithm uses Kernel Density Estimation (KDE) to transform the spatial mRNA coordinates into gene expression probability densities that are subsequently cell typed and then projected into the final image of the cell-type map. SSAM-lite is an integrated pipeline aimed at simplifying exploratory data analyses of SRT data with only a few clicks in a web browser. The pipeline workflow combines state-of-the-art web programming libraries such as *Bootstrap*, *plotly.js*, and *TensorFlow.js* (Figure 1A). The modern web interface with convenient interactive elements was generated using the *Bootstrap* library, which provides a large body of CSS functions for creating a state-of-the-art and user-friendly layout. In particular, the layout scripts for SSAM-lite make use of *Bootstrap*’s sophisticated scalable grid layout that optimizes user experience on a range of devices from handhelds to desktop machines. The data preparation and presentation routines were implemented using *plotly.js*, and *TensorFlow.js* was chosen to implement a machine learning backend.

**Figure 1.**
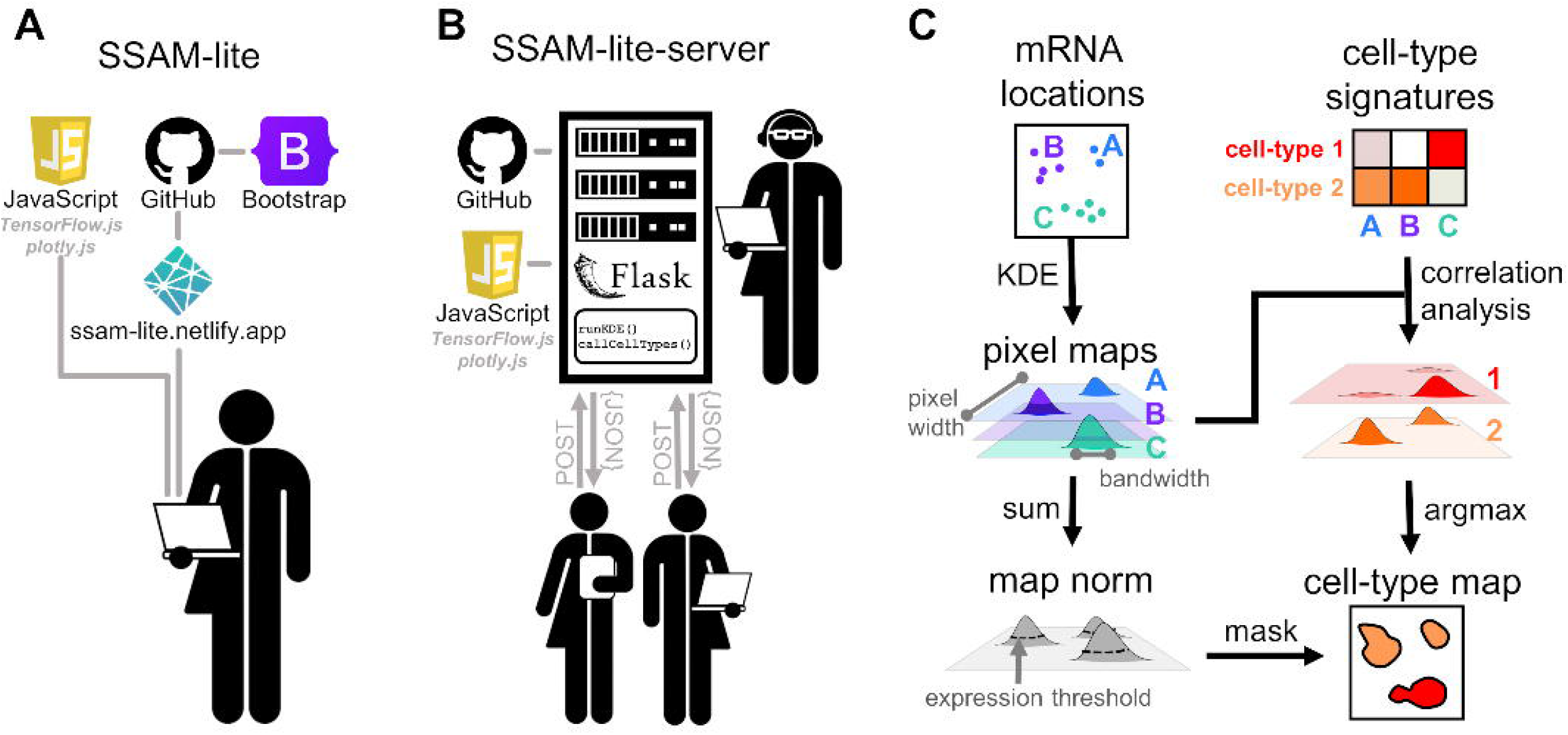
schematic of SSAM-lite. (A) Schematic diagram of SSAM-lite, accessible as a web browser application, and (B) a locally installed SSAM-lite-server. (C) Schematic of the underlying data processing algorithm proposed by SSAM.

A typical SSAM-lite workflow can be summarized in three steps: data upload, parameter selection, and optimization, and the final analysis phase. Each step has a dedicated area in the web interface (Figure 2).

**Figure 2:**
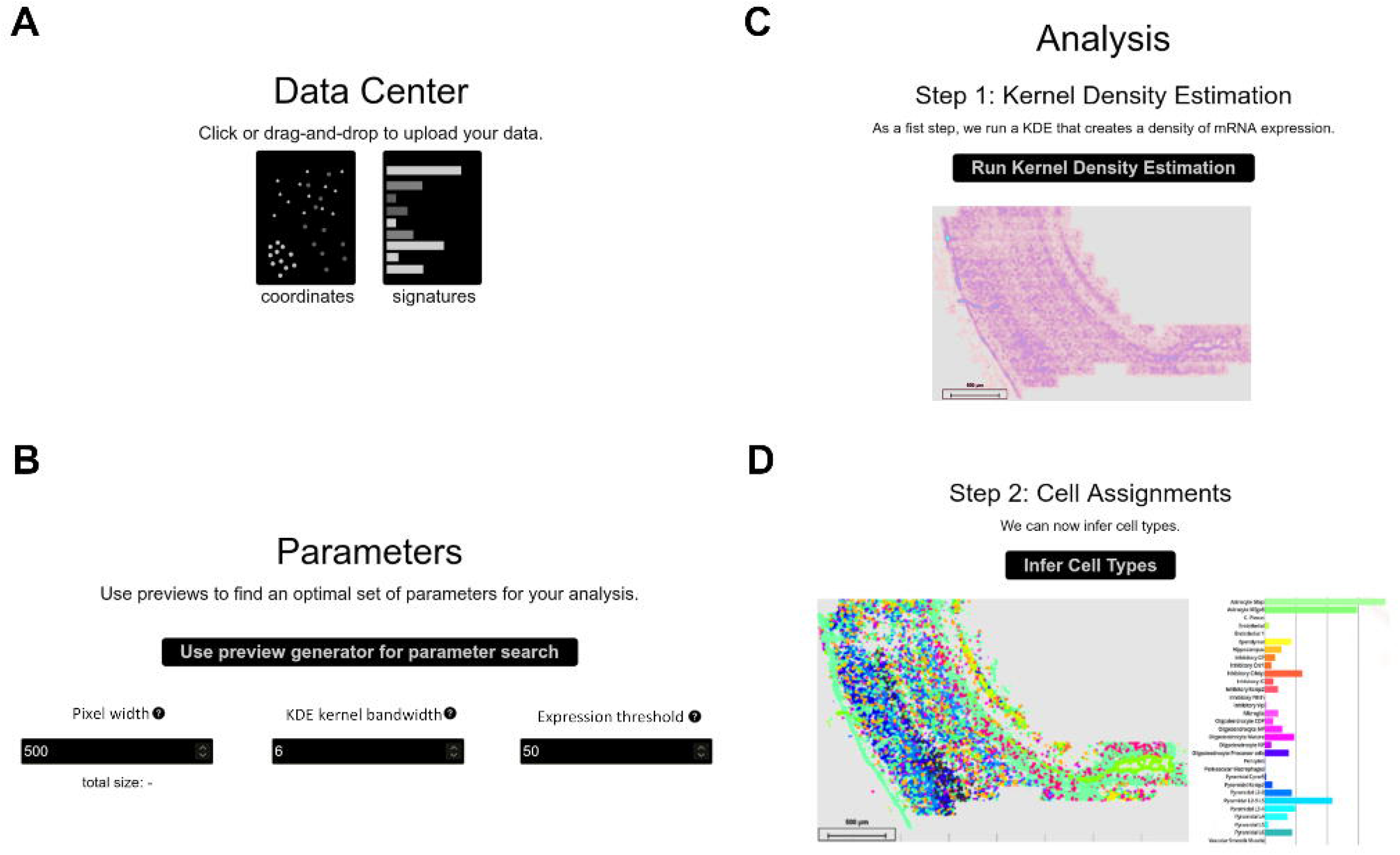
the SSAM-lite interface. The panels display the sections of the SSAM-lite web page demonstrated on osmFISH data of the mouse SSp (Codeluppi et al., 2018): (A) the data center for uploading data; (B) the parameters selection and optimization section; (C) the first analysis section for displaying the results of the KDE analysis; and (D) the second analysis section for displaying the final cell-type map image.

#### 3.1.1 Data upload

Data upload is performed in the *Data Center* section by either using drag-and-drop or an interactive file selection window (Figure 2A). The user needs to provide a file with mRNA coordinates from an SRT experiment alongside a so-called signature file that contains gene expression signatures for the cell types of the tissue of interest. Both input files are plain text csv files. The mRNA coordinate file contains gene names and the x- and y-coordinates of all molecules in the analyzed image, consistent with the *DecodedSpot* format defined by the Starfish pipeline (http://github.com/spacetx/starfish). The signature file contains a gene expression matrix with cell types as rows and gene names as columns. The values can either be binary or be normalized gene expression.

After loading, the mRNA molecule coordinate data is displayed in an interactive scatter plot using *plotly.js*’s *scattergl* layout, which is designed explicitly to handle large data sets. The plot is designed to be interactive, so the user can zoom in to investigate local mRNA expression or hide parts of the data to reveal the expression patterns of individual genes. The expression signature matrix is also displayed in an interactive plot after loading using *plotly.js*’s *heatmap* layout, which provides an overview of the data through color coding and by displaying hovering information on each gene-cell type expression indicator.

#### 3.1.2 Parameter selection and optimization

In the *parameter selection and optimization* section, the user can interactively tune the input parameters for the SSAM spatial modeling algorithm (Figures 2B, 3). The three most important parameters of the SSAM algorithm are the *bandwidth* of the Gaussian KDE function, the *pixel width* of the output cell-type map, and the total *expression threshold* value. The *bandwidth* parameter is necessary to accurately model the local spatial molecular dynamics. To model expression in a sparse dataset (e.g. 3-5 mRNA molecules per cell) a larger bandwidth would need to be employed, and in a dense dataset (e.g. 20-30 mRNA per cell) a smaller bandwidth should be sufficient. As a guideline, we suggest using values between 2 and 25 micrometers based on analysis of dense and sparse datasets (Figure 4). The *pixel width* of the cell-type map determines the memory footprint and the accuracy of the internal spatial gene expression model. The *expression threshold* parameter defines the gene expression signal threshold for the foreground (i.e. parts of the image with high gene expression, likely originating from cells) and background (i.e. parts of the image with low gene expression), hence discerning actual spatial expression patterns from background noise. A high number of extracellular, diffused mRNA spots requires a higher expression threshold, where the optimal value differs greatly across data sets.

**Figure 3.**
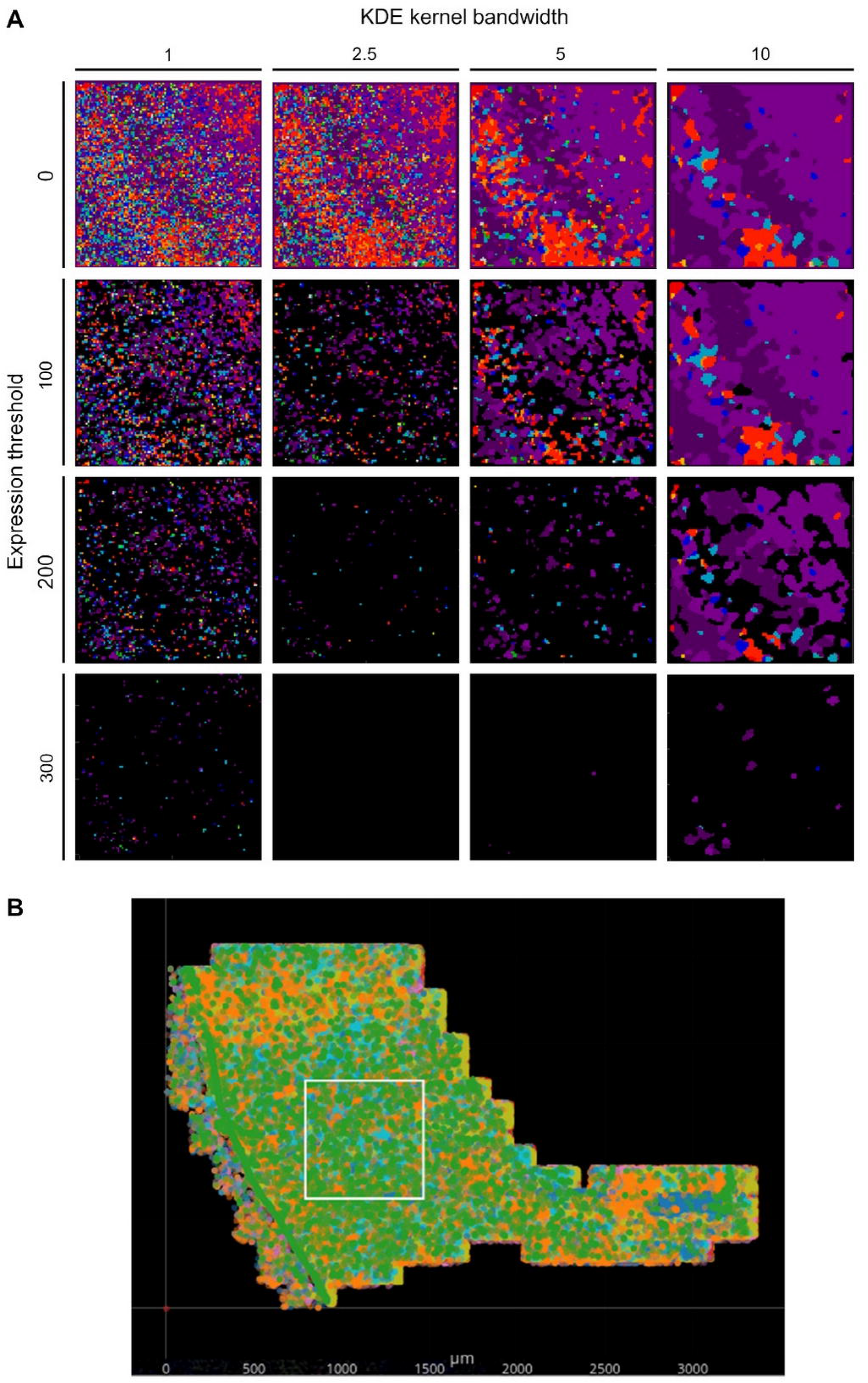
Effects of the expression threshold and bandwidth parameters on the cell-type map. The entire SSAM-lite pipeline with different input parameters (A) was applied to the square subsection of the osmFISH mouse SSp dataset highlighted in (B). (A) Shows the output cell-type maps for different combinations of expression thresholds (0, 100, 200, 300) as rows and bandwidth parameters (0, 2.5, 5, 10) as columns. The effect of the bandwidth and expression threshold parameters are discussed in the Methods section.

**Figure 4:**
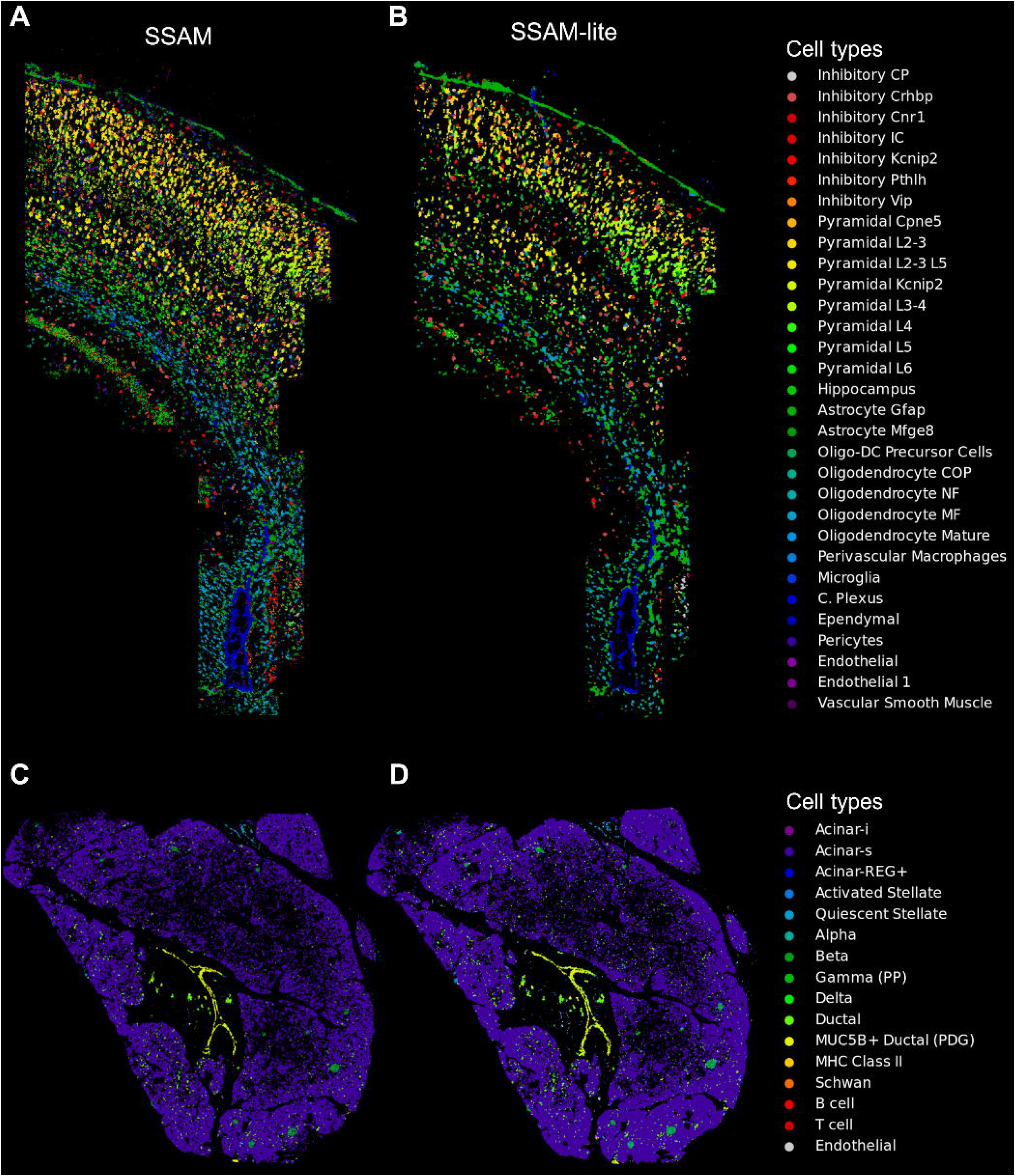
SSAM-lite generates accurate cell-type maps. Demonstrative cell-type maps for osmFISH data of the mouse SSp generated by (A) SSAM and (B) SSAM-lite, and ISS data of human pancreas generated by (C) SSAM and (D) SSAM-lite. Resultant cell-type maps generated by SSAM are similar to previous publications (Park et al., 2021; Tosti et al., 2021). Cell-type colors of the original SSAM figures were modified to match the SSAM-lite figure.

These parameters can be set in numerical input fields and the analysis of the full data set can be started. However, the user can also try to optimize the parameters on a small section of the image before starting the complete image analysis. This will launch an initial small-scale analysis with instant output to the screen and will show three figure panels that allow for direct evaluation of the chosen parameters.

Of these panels, the left figure panel is an interactive *plotly.js scattergl* plot of the entire mRNA location data set, which can be used to define the local sub-section of the overall data set that is used for optimization. A rectangle displays the currently chosen sub-section, and the location of the subsection can be changed interactively by clicking onto the desired new central spot in the scatterplot.

The middle figure panel shows an intermediate output of the KDE for the chosen sub-section from the first subfigure using *plotly.js*’s *heatmap* layout. The heatmap is a spatial representation of SSAM-lite’s internal model of integrated local signal strength, with the heatmap value indicating the probability for the presence of a cell at a particular location. The value of the modeled signal for each pixel is color-coded and shows up when hovering over it with the mouse pointer. The heatmap is especially useful for choosing an appropriate *expression threshold* parameter from the signal strength landscape. The KDE figure panel also provides a visual impression of the amount of smoothing produced by the KDE, which helps the user to set the *bandwidth* parameter. The *bandwidth* parameter should be large enough to smooth out noise and integrate mRNA signals belonging to the same spatial structure, but low enough to keep individual spatial structures separate and retain their shape. The heatmap plot gets updated in real-time whenever the subsample location or KDE parameters change, and in practice, the parameters can be set reasonably after 2-3 trials.

The rightmost figure panel shows the final output cell-type map of the SSAM-lite algorithm for the chosen tissue subsection. The cell-type map is useful to identify persisting noise in the output, which can be reduced by adjusting the *bandwidth* and/or the *expression threshold* parameters. The cell-type map panel is updated whenever the parameters change.

#### 3.1.3 Analysis and visualization

The last section is dedicated to data analysis and the visualization of results. The section provides interfaces to the two more resource-intensive *TensorFlow.js* backend functions that perform the KDE and the correlation analysis.

##### 3.1.3.1 Kernel Density Estimation (KDE)

Once the parameters are optimized, the user can perform the KDE, which typically takes below a minute to generate SSAM-lite’s internal, pixel-based spatial model of local signal strength (Figure 2C). In a pre-processing step, the mRNA coordinates are rescaled linearly to fit the user-defined *pixel width* of the spatial model. The respective height is determined to match the vertical spread of the coordinate data and the bandwidth parameter is scaled accordingly to match the new internal unit of computation. SSAM-lite computes an independent local signal strength pixel matrix for each type of mRNA defined in the input data. For this, a large *TensorFlow.js* buffer is initiated by stacking all empty pixel matrices. The KDE implementation in SSAM-lite employs two heuristics to optimize computing performance. The first is to iterate over all mRNA locations and round them to their closest output pixel, allowing us to use a pre-calculated Gaussian mass function for all mRNA spots. The second is to ignore long tails of the Gaussian mass function by limiting its calculation to two bandwidths.

After KDE computation is completed, the collected sum of all pixel matrices is displayed using a *plotly.js heatmap* layout analogous to the optimization panel. If the results do not match expectations, parameters can be adapted and the KDE function can be re-run. Otherwise, the user can move on to generate the cell-type map.

##### 3.1.3.2 Correlation Analysis and Cell-Type Map Generation

As in the original SSAM algorithm, the last step of analysis computes the cell-type map through correlation analysis with known gene expression signatures (Figure 2D). The combined expression arrays of each x- and y-location in the stacked pixel matrixes are compared to the expression signature data and each pixel is assigned the cell type with the highest correlating signature. All pixels whose sum across matrices are below the user-defined *expression threshold* parameter are considered background and not assigned any cell type. The final result is displayed as a cell-type map using a modified version of *plotly.js*’s *heatmap* layout. The heatmap element is fed with a custom generated list of colors and altered to display the x- and y-coordinates and the assigned cell-type name during mouse hover events. The plot offers *plotly.js*’s elementary functions like zooming, panning, resetting as well as a save to disk option. Furthermore, a custom scale bar is added that adapts to the current zoom factor and displays the bar width in micrometers.

##### 3.1.3.3 Cell-type localization and abundance

An important part of the downstream analysis of the cell-type map is the localization of cell types and the quantification of cell-type signals in the entire and parts of the tissue (Figure 2D, 5A). We therefore implement an interactive *barplot* that quantifies the relative cell-type abundance based on classified pixels in the current view of the cell-type map. This quantification is updated when zooming into or panning over different regions (Figure 5B, 5C). The user can also provide custom color palettes and select only certain cell-types to be rendered by double-clicking the cell-type labels (Figure 6).

**Figure 5.**
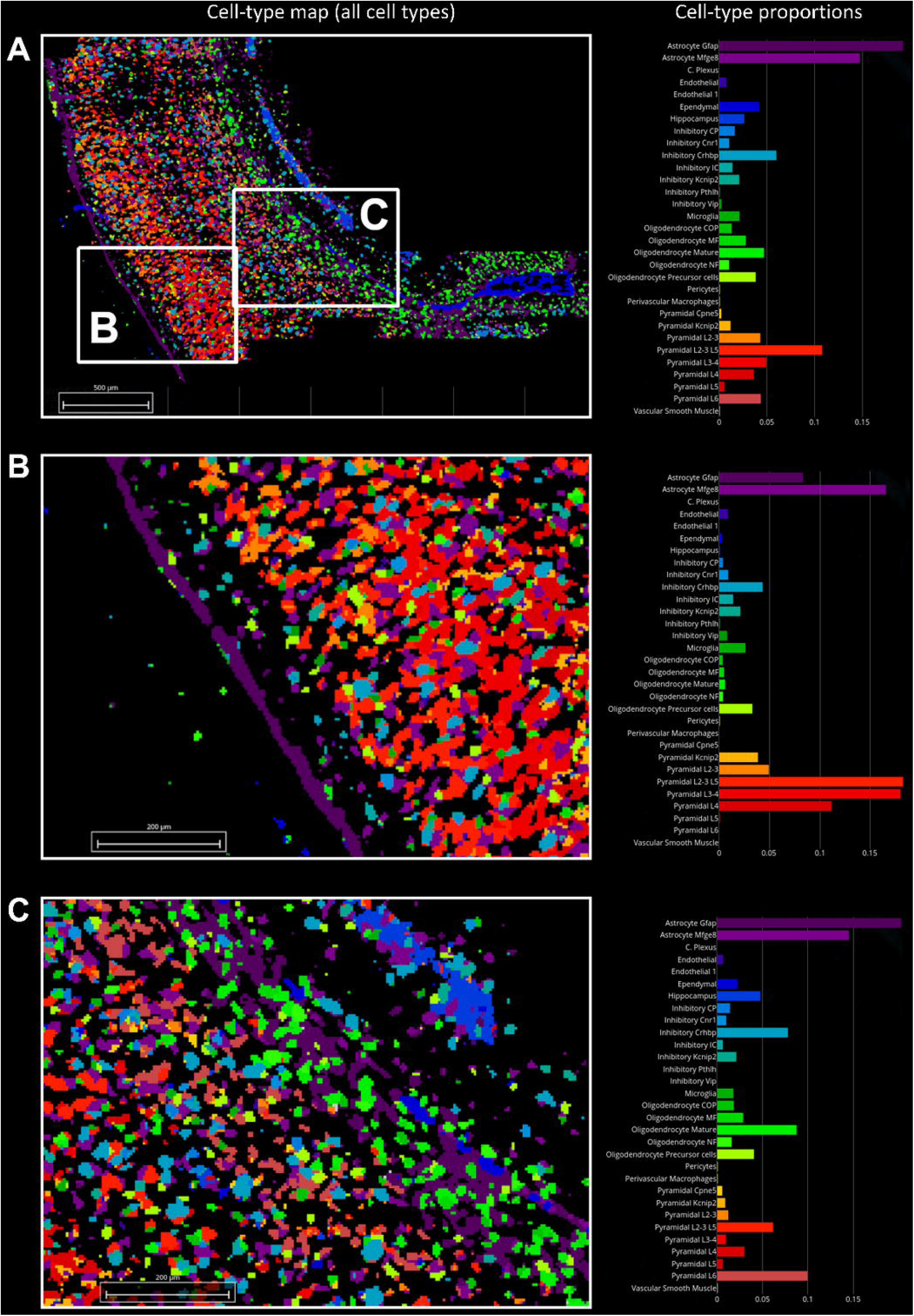
Regional cell-type proportion estimates. Cell-type map and relative cell-type proportion estimates in the (A) entire mouse SSp, (B) zoom-in of the pia and cortex layers 1-4, and (C) zoom-in of cortex layers 4-6 and the hippocampus. Panel B shows enrichment of L2-3, L2-3 L5, L3-4 and L4 pyramidal cell types, and panel C shows enrichment of L2-3 L5, L4, and L6 cell types.

**Figure 6.**
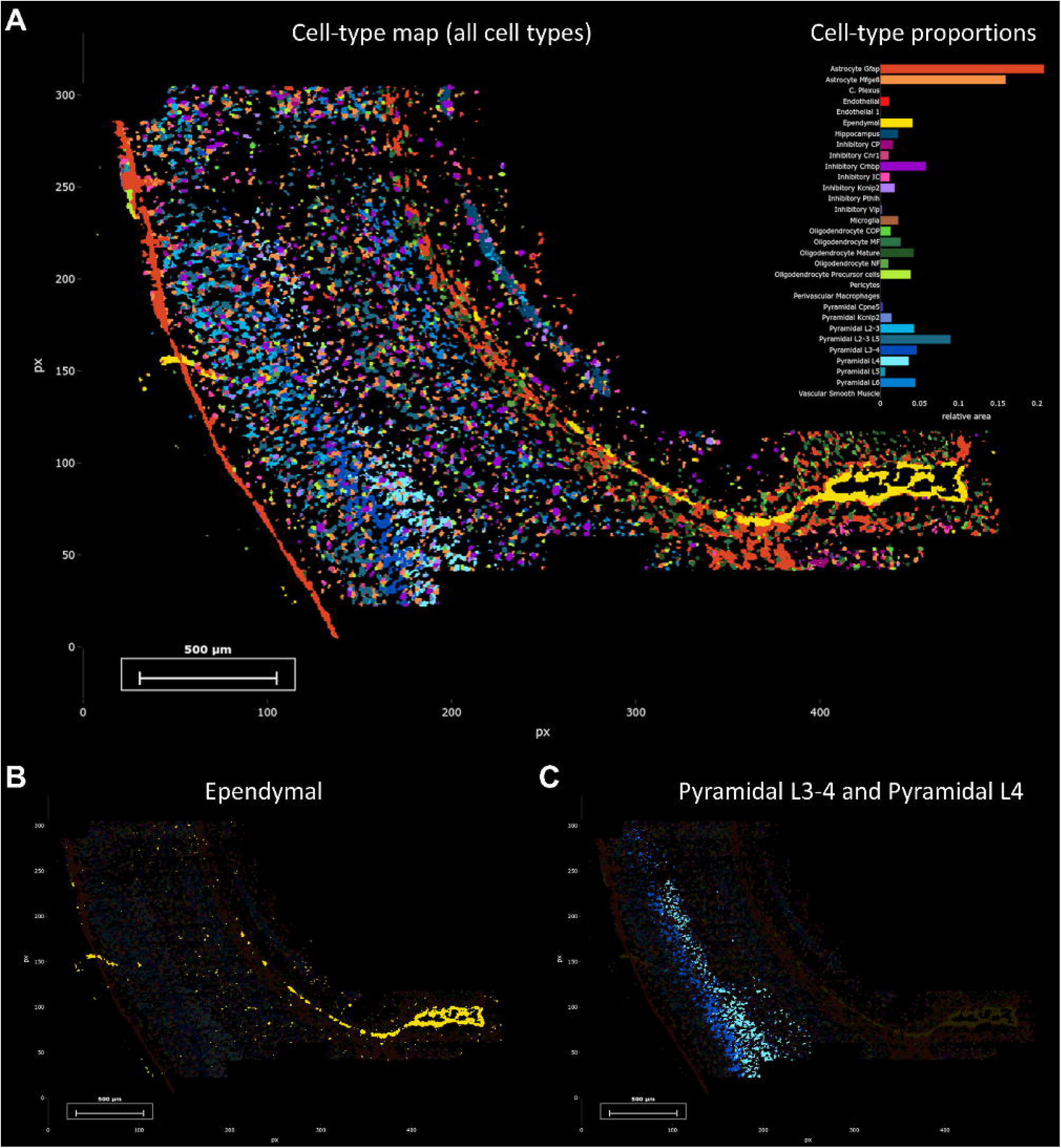
Custom color palettes and cell-type highlighting. Cell-type map of (A) all cell types in the mouse SSp, (B) only ependymal, (C) and pyramidal L3-4 and pyramidal L4 cell-types. The color palette shows cell types rendered using the same colors as in Codeluppi et al (https://doi.org/10.1038/s41592-018-0175-z).

The code itself is documented and organized according to the model-view-controller paradigm, which allows the user to easily adapt the code base to the needs of their own specific project. One example would be to use an alternative kernel shape, e.g. a circular Epanechnikov kernel could be achieved by adding a logical threshold expression to the *runKDE* function inside *model.js*. Any changes are integrated into the code execution right away and available after a simple browser page refresh.

### 3.1.4 SSAM-lite-server

SSAM-lite is an efficient tool that is dependent on client-side hardware. While we demonstrate that a modest laptop is capable of processing real-world SRT datasets (Figure 5), we also recognize possible limitations due to client-side hardware constraints. To address this issue, we developed a server-side version called SSAM-lite-server (Figure 1A). SSAM-lite-server runs the computationally expensive KDE and cell assignment algorithms at the server-side. SSAM-lite-server preserves the overall implementation of SSAM-lite in Javascript, HTML, CSS, and allows a server running a *Flask* (v0.8) framework to take over computationally expensive functionalities of SSAM-lite. *Flask* was chosen due to its lightweight nature and extensibility. To further make the backend data structures memory-efficient we use Python’s numerical library *NumPy* (v1.20.3). Python’s *pandas* package (v1.3.2) is used to handle the signature data. For privacy preservation, the data streamed to the server for processing does not persist on the server file systems but is only stored in memory for the duration of the computation.

SSAM-lite-server runs the KDE algorithm by streaming variables such as coordinates, signature matrix, input and output image width, bandwidth, gene expression threshold to the server as an Ajax POST request, which then returns JSON objects to the user. The server-side computation includes the computation of KDE and the generation of the cell-type map.

To enhance the overall security, SSAM-lite-server offers the option to host all libraries locally, thus enabling SSAM-lite-server to run in closed networks without an internet connection.

### 3.1.5 Benchmarking

Benchmarking was carried out on a Lenovo X1 Carbon laptop with Intel Core i7-8565u CPU, 16GB of RAM, and Windows 10. We used Google Chrome (v93.0) to run SSAM-lite (v0.1.0). The benchmark was performed using the Chrome DevTools Performance monitor to evaluate the runtime of the *runFullKDE* function and the maximum memory heap while carrying out a complete analysis (not using the parameter preview) with the pixel width of the cell-type map set to 500, the kernel bandwidth to five and the expression threshold for assigning cell types to two.

To simulate different complexities of the mouse brain primary somatosensory cortex (SSp) data we performed downscaling and upscaling of the data. A 0.5x dataset was created by randomly downsampling to 50% of the molecules present in the coordinate file. A 2x data set was created by appending the mRNA coordinate locations to itself after carrying out a pixel shift of 1 micrometer along both axes to each of the molecules. A 3x dataset was created by pixel shifting the original coordinate matrix by −1 micrometer and appending it to the 2x coordinate matrix. Finally, the 5x dataset was created by appending the dataset to itself, the first time pixel-shifting +1 along x and y, the second time +2, and so on. Each of the above datasets was then tested in 3 replicates.

## 4 Results

To demonstrate equivalent cell-type map performance to our previously published SSAM algorithm, we applied SSAM-lite to 2 datasets using a laptop computer (see Benchmarking section in Methods). The first dataset was mouse SSp profiled by osmFISH (Zeisel et al., 2015; Marques et al., 2016), profiling 1,802,589 mRNA spots for 33 genes and 31 cell-types signatures derived from scRNAseq (Zeisel et al., 2015; Marques et al., 2016). The coordinate matrix was uploaded and rendered in 4 seconds on average, and the uploading and rendering time for the signature matrix was negligible in comparison. The cell-type map width was set to 1500, KDE bandwidth to 2.5, and the gene expression threshold to 13. The resultant image of the cell-type map was very similar to those previously published (Figure 3A, 3B). To demonstrate SSAM-lite’s performance on a sparse dataset, we applied it to human pancreas profiled by ISS, profiling 461,078 mRNA spots for 138 genes and 16 cell-type signatures (Tosti et al., 2021). The cell-type map width was set to 750, KDE bandwidth to 22, and the gene expression threshold to 2.4. The resultant image of the cell-type map was highly comparable to those previously published (Figure 5C, 5D).

To investigate how SSAM-lite performance scales with regards to memory requirements and CPU time, we performed a synthetic benchmark on the mouse brain SSp dataset with different dataset sizes (Figure 7). Overall, the CPU time for calculating the KDE (Figure 7A) scales linearly with the number of profiled mRNA molecules. Further, the total memory footprint for a complete analysis also depends linearly on the dataset size (Figure 7B).

**Figure 7.**
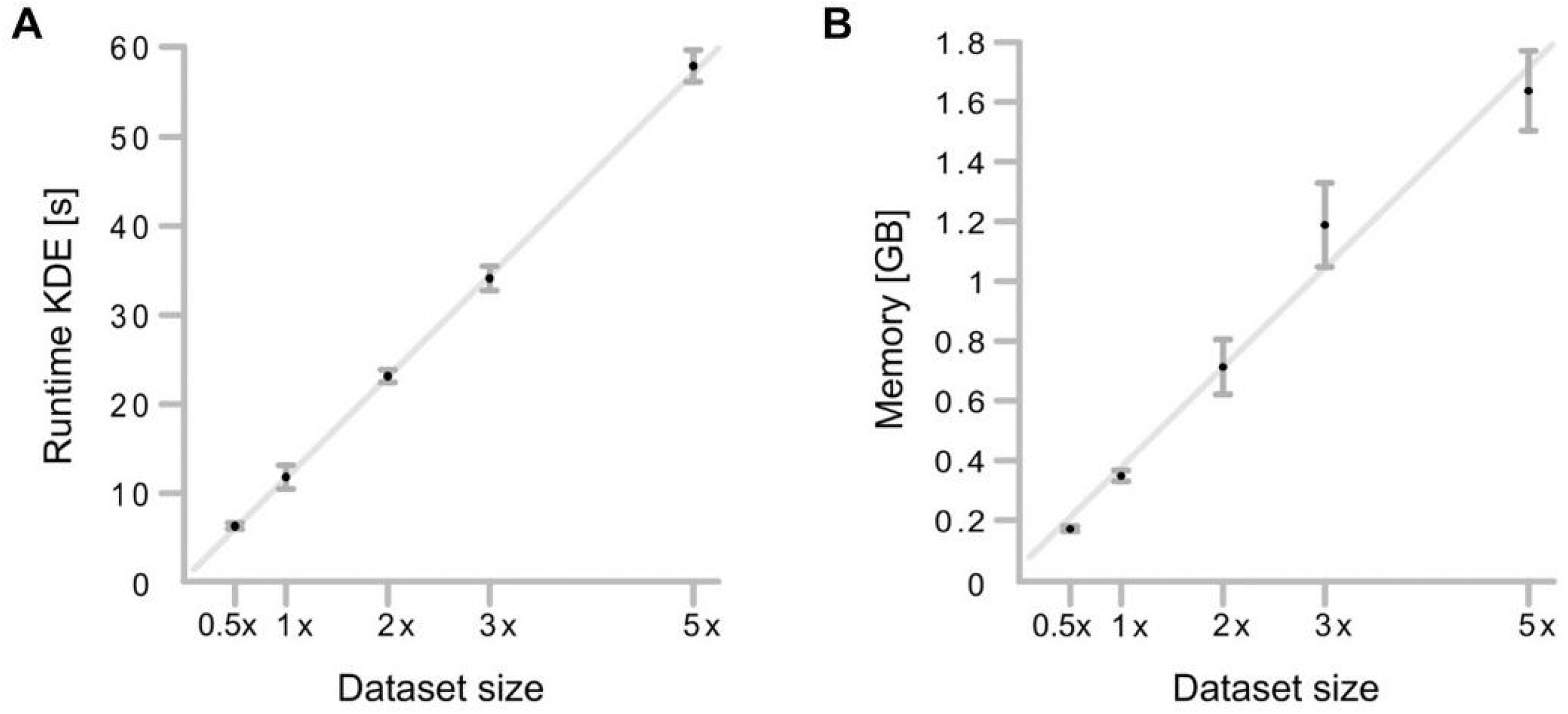
SSAM-lite runtime and memory usage scale linearly with dataset size. The simulation was based on the mouse SSP osmFISH dataset which has 1.8 million mRNA spots. (A) Computation time for the KDE step scales linearly with data set size (R^2^ = 0.9996, F-value=9445 and p-value < 2.41e-6). (B) Maximum RAM consumption scales linearly with dataset size (R^2^=0.9738, F-value=149.9 and p-value < 1.18e-3). The SSP data set was down-sampled by 50% and up-scaled by a maximum factor of 5. Black dots are mean values. Error bars are standard deviations. (n=3). Grey line is a linear fit. The 0.5x dataset took on an average 6.32 seconds to run with a memory footprint of 171 MB. In comparison, the whole dataset took on average 11.81 seconds with a memory footprint of 343 MB. Whereas the largest 5x dataset on average took 57 seconds with a memory footprint of 1.6 GB. These results suggest that there is a linear increase of memory and CPU requirements with an increasing number of mRNA molecules profiled in a single experiment.

## 5 Discussion

Analysis of spatial transcriptomics data was so far limited by excessive hardware requirements and an understanding of navigation in a terminal window using the Linux command line. With SSAM-lite we overcome these limitations by providing an easy-to-use graphical user interface that runs in any modern web browser on common laptop computers. Input files are text files that can be loaded by drag-and-drop into the browser window. This circumvents the need to provide certain command-line arguments or editing of configuration files. SSAM-lite makes the analysis of spatial transcriptomics data accessible to a broad range of researchers that may not have a high-performance computing cluster or experience with command-line tools. SSAM-lite was able to generate similar results to those previously published(Park et al., 2021; Tosti et al., 2021) in only a few minutes. SSAM-lite provides an easy-to-use interface to analyze high-dimensional SRT data to the wider biomedical research community. In addition, we see the additional utility in SSAM-lite for SRT data generators to perform rapid quality control of experiments and to provide customers with an easy-to-use exploratory tool. We also expect that specialized computational scientists may want to use SSAM-lite to rapidly identify optimal parameters for downstream analysis and to compare the resultant cell-type map of more parameterized and resource-hungry analysis tools.

In addition, SSAM-lite-server mitigates much of the computational burden to the server-side, enabling analysis of very large datasets, and also analysis of datasets on mobile devices. The stand-alone implementation of SSAM-lite-server is amenable to networks with limited access to the internet such as in many university hospitals.

## 7 Author Contributions

ST and NI conceived and designed the study. ST programmed the SSAM-lite and SSAM-lite-server software. SS made programming contributions to SSAM-lite-server. ST, SDM, and NI wrote the manuscript. SS, NMB tested and documented the software, made programming contributions to SSAM-lite, wrote the user guide, and revised the manuscript. RE proofread and corrected the manuscript. All authors contributed to the article and approved the submitted version.

## 9 Acknowledgments

We would like to thank Jeongbin Park and Wonyl Choi for conceiving the idea of SSAM.

## 10 Data Availability Statement

Publicly available datasets were analyzed in this study. This data can be found here: https://zenodo.org/record/5517607.

